# Theta oscillations in anterior cingulate cortex and orbitofrontal cortex differentially modulate accuracy and speed in flexible reward learning

**DOI:** 10.1101/2021.09.14.460307

**Authors:** Tony Ye, Juan Luis Romero-Sosa, Anne Rickard, Claudia G. Aguirre, Andrew M. Wikenheiser, Hugh T. Blair, Alicia Izquierdo

## Abstract

Flexible reward learning relies on frontal cortex, with substantial evidence indicating that anterior cingulate cortex (ACC) and orbitofrontal cortex (OFC) subregions play important roles. Recent studies in both rat and macaque suggest theta oscillations (5-10 Hz) may be a spectral signature that coordinates this learning. However, network-level interactions between ACC and OFC in flexible learning remain unclear. We investigated the learning of stimulus-reward associations using a combination of simultaneous *in-vivo* electrophysiology in dorsal ACC and ventral OFC, partnered with bilateral inhibitory DREADDs in ACC. In freely-behaving male and female rats and using a within-subject design, we examined accuracy and speed of response across distinct and precisely-defined trial epochs during initial visual discrimination learning and subsequent reversal of stimulus-reward contingencies. Following ACC inhibition there was a propensity for random responding in early reversal learning, with correct vs. incorrect trials distinguished only from OFC, not ACC, theta power differences in the reversal phase. ACC inhibition also hastened incorrect choices during reversal. This same pattern of change in accuracy and speed was not observed in viral control animals. Thus, characteristics of impaired reversal learning following ACC inhibition are poor deliberation and weak theta signaling of accuracy in this region. The present results also point to OFC theta oscillations as a prominent feature of reversal learning, unperturbed by ACC inhibition.

## INTRODUCTION

Both humans and nonhumans alike make a multitude of choices based on adaptations to their environment in order to maximize rewards while minimizing losses. This type of flexible reward learning is thought to rely on the prefrontal cortex. Substantial evidence in primates indicates that anterior cingulate cortex (ACC) and orbitofrontal cortex (OFC) have essential roles in learning about actions and stimuli, respectively (Rudebeck, Behrens et al. 2008, Camille, Tsuchida et al. 2011, Luk and Wallis 2013). Though primate and rodent ACC and OFC are not equivalent structures, we may still learn fundamental principles of their functions in the rodent (Rudebeck and Izquierdo 2021). A mechanistic investigation of their interaction in stimulus-based learning is missing in the broader literature.

Lesion or inhibition of OFC impairs multiple forms of reward learning and in rats this has most extensively been observed in the reversal phase (Chudasama and Robbins 2003, Izquierdo, Darling et al. 2013), particularly in spatial- or action-based reversal learning (Boulougouris, Dalley et al. 2007, Riceberg and Shapiro 2012, Dalton, Wang et al. 2016, Groman, Keistler et al. 2019). Classic recording studies in rat OFC during (olfactory cue) reversal learning (Schoenbaum, Chiba et al. 1999) show that neural activity represents an integration of the motivational significance of a reward and its specific predictive cue, critical for reversal but not discrimination learning (Schoenbaum, Chiba et al. 1999, Marquardt, Sigdel et al. 2017). Conversely, lesion or inhibition of ACC results in decrements in (visual) discrimination performance (Chudasama, Passetti et al. 2003), but has no effect on fully-predictive, deterministic reversal learning (Bussey, Muir et al. 1997, Schweimer and Hauber 2006). Yet ACC neurons in rats integrate motor-related information with expectations of future outcomes (Cowen, Davis et al. 2012, Hyman, Holroyd et al. 2017), encode choice value (Mashhoori, Hashemnia et al. 2018) and maintain trial-by-trial performance (Akam, Rodrigues-Vaz et al. 2021), all processes that would be vital not just to action-based learning but also to stimulus-based reversal learning. This growing body of evidence supports a general role for ACC in maintaining a model about the reward environment and making predictions about future choices associated with either actions or stimuli.

More importantly, how these two regions interact is unclear. Several groups have recorded single-unit and local-field activity simultaneously in both areas and uncovered overlapping functions, in contrast to the interference studies (Rudebeck, Walton et al. 2006, Rudebeck, Behrens et al. 2008). For example, there is reported evidence of ACC-OFC task-related coherence in theta to low beta (4-20 Hz) oscillations as rats respond to or choose a higher-valued reward, suggesting a functional connectivity underlying relative reward value (Fatahi, Haghparast et al. 2018, Fatahi, Ghorbani et al. 2020, Amarante and Laubach 2021). It is therefore conceivable that these two regions work in a complementary manner for reward-based learning and choice (Hunt and Hayden 2017, Hunt, Malalasekera et al. 2018) potentially with theta oscillations in OFC coordinating reward signals (Marquardt, Sigdel et al. 2017, Knudsen and Wallis 2020) with ACC (Fatahi, Haghparast et al. 2018). However, network-level interactions and the spectral signatures in flexible reward learning remain unclear.

Given that ACC is densely innervated by regions involved not only in evaluating options but also preparing and executing actions, we hypothesized that ACC theta modulates overall learning of stimuli. To test this, we investigated learning using a combination of simultaneous *in-vivo* electrophysiology in dorsal ACC (dorsal area 32/24, van Heukelum, Mars et al. (2020)) and ventral OFC, partnered with inhibitory Designer Receptors Exclusively Activated by Designer Drugs (DREADDs) in ACC. We recorded from posterior VO as it is richly interconnected with ACC (Hoover and Vertes 2011, Barreiros, Panayi et al. 2021). Additionally, a thorough consideration of both accuracy and speed of response during learning may be particularly informative when identifying neural signatures of flexible learning (Aguirre, Stolyarova et al. 2020, Harris, Aguirre et al. 2021), so we include analysis of both here.

We found that ACC inhibition during the discrimination phase had no functional impact on initial performance. In contrast, ACC inhibition in early reversal learning promoted a random response strategy, disrupted ACC signaling of correct vs. incorrect trials, and hastened speed of incorrect choices. The same pattern was not observed in viral control animals. Though ACC inhibition produced these changes in accuracy and speed in the reversal phase, it did not affect OFC theta signaling of accuracy in reversal learning, suggesting an independent process in OFC.

## RESULTS

### ACC inhibition in early, not late, reversal learning promotes random responding

Fifteen rats were prepared with either inhibitory (hM4Di) DREADDs (n=8, 4 females) or enhanced Green Fluorescent Protein (eGFP) null virus (n=7, 3 females) on putative projection neurons in ACC. Eight of these animals (n=4 hM4Di, n=4 eGFP) were also implanted with custom-constructed 16-channel fixed electrode arrays unilaterally in ACC and OFC, in the same surgery as virus infusion. All rats were tested using a within-subject design for drug administration (**Figure 1**). Rats were administered clozapine-N-oxide (CNO) once they reached criterion-level performance (80%) on the initial discrimination learning phase to assess the impact of ACC inhibition on mastery-level performance (Bussey, Muir et al. 1997, Chudasama, Passetti et al. 2003). Subsequently in the reversal phase, they were administered CNO or vehicle (VEH) in counterbalanced order: CNO administered early in learning (to 50% correct) then switched to VEH until 80%, or vice versa (VEH to 50%, then CNO to 80%). We used a CNO dose for ACC inhibition that has previously resulted in behavioral effects on two different reward learning tasks (Stolyarova, Rakhshan et al. 2019, Hart, Blair et al. 2020), and within the dose-range exerting the least off-target effects (Jendryka, Palchaudhuri et al. 2019).

**Figure 1.**
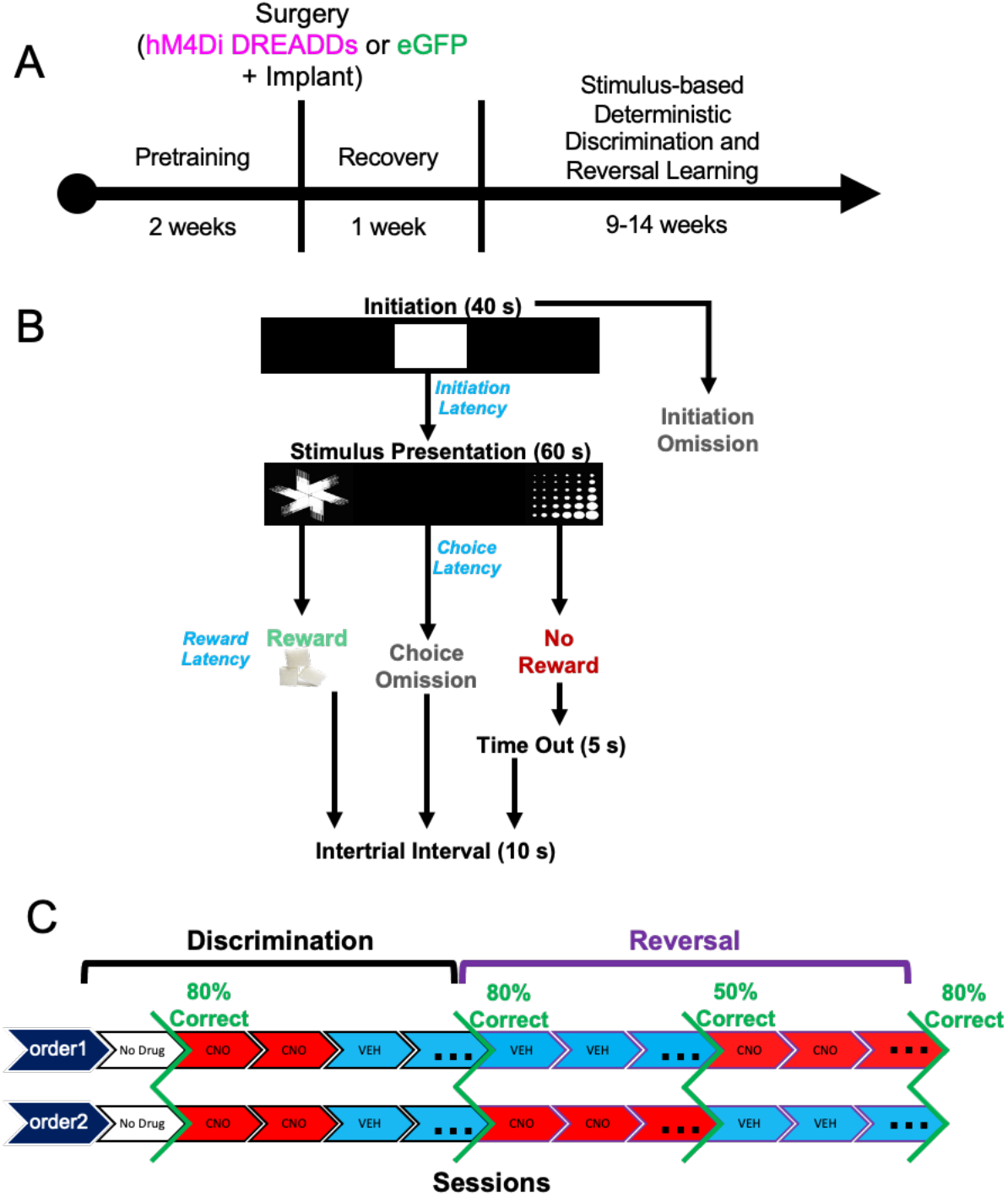
Experimental timeline, task structure, and drug administration. (**A**) After approximately 2 weeks of pretraining, rats underwent bilateral viral infusion of inhibitory hM4Di DREADDs or null virus enhanced Green Fluorescent Protein (eGFP) in ACC along with chronic implantation of electrode arrays targeting both ACC and OFC unilaterally in each animal. (**B**) Following a 1-week recovery period, rats commenced the stimulus-based visual discrimination and reversal learning task, with rewarded-stimulus assignment (e.g., “fan” vs. “marbles”) counterbalanced across rats. (**C**) Drug administration (clozapine-N-oxide, CNO or vehicle, VEH) was administered once criterion was reached for initial discrimination to probe the effect of ACC inhibition on mastery-level discrimination performance. Subsequently, in the reversal phase, the order of drug administration was counterbalanced across animals.

There was no significant difference between hM4Di and eGFP animals in initial discrimination learning, though two rats from each virus group failed to learn (sessions to criterion: t(11)= −0.250, *p*=0.807), **Figure 2A**. A mixed-effects GLM was used to analyze probability correct, with drug, virus, and sex as between-subject factors, session as a within-subject factor, and individual rat as random factor. There was no functional consequence of ACC inhibition on mastery level performance (i.e., no significant effect of drug, virus, or interaction of drug x virus in a mixed-effects GLM; GLM formula: *γ* ∼ *[1 + Drug*\**Session*\**Virus*\**Sex + (1 + Session*| *Rat)*], **Figure 2B**. A significant effect of session, and interactions of session x virus, and session x virus x drug were observed, **Table 1**. When post-hoc Bonferroni post-hoc comparisons were conducted, we found eGFP (p<0.01) but not hM4Di (p=0.11) groups decreased in probability across session, though VEH performance was greater than CNO in both eGFP and hM4Di groups (both p<0.01s).

In contrast to the acquisition curves that demonstrated mastery of the initial visual discrimination, all animals exhibited difficulty learning reversals, rarely achieving above 60% after 10 sessions (**Figure 2C**), similar to a recent report (Harris, Aguirre et al. 2021). A mixed-effects GLM was also used to analyze reversal learning. This analysis resulted in a significant drug x virus interaction (β_drug x virus_ = 0.258, t(178) = 2.56, *p*=0.01). Post-hoc Bonferroni-corrected tests revealed a nonsignificant effect of drug in the hM4Di group (*p*=0.07) and in the eGFP group (*p*=1.0). There were other interactions with sex and session that did not result in significant effects with Bonferroni-corrected post-hoc follow-up (**Table 2**). However, when we added drug order to the GLM model we found a significant drug x drug order interaction (β_drug x drug order_ = −0.56983, t(162) = −3.39, *p*<0.001) with an effect of drug only in the hM4Di group (post-hoc Bonferroni-corrected tests: p=0.01 for hM4Di, p=0.68 for eGFP). Early inhibition of ACC in the reversal phase brought performance to chance and maintained it there (**Figure 2D**), in contrast to rats given VEH first which instead showed the expected tendency to follow the previous stimulus-reward assignment on early reversal sessions during VEH (i.e., more perseveration), but reached higher accuracy levels later, even when administered CNO. Interestingly, CNO had the opposite effect in eGFP animals: when administered early, rats exhibited the expected learning trajectory, but when administered CNO at 50% accuracy, performance remained at chance level (**Figure 2C**). Collectively, the pattern of results reveals a differential effect of CNO in hM4Di and eGFP animals, with ACC inhibition producing a random response strategy early in reversal learning.

**Table 1.**
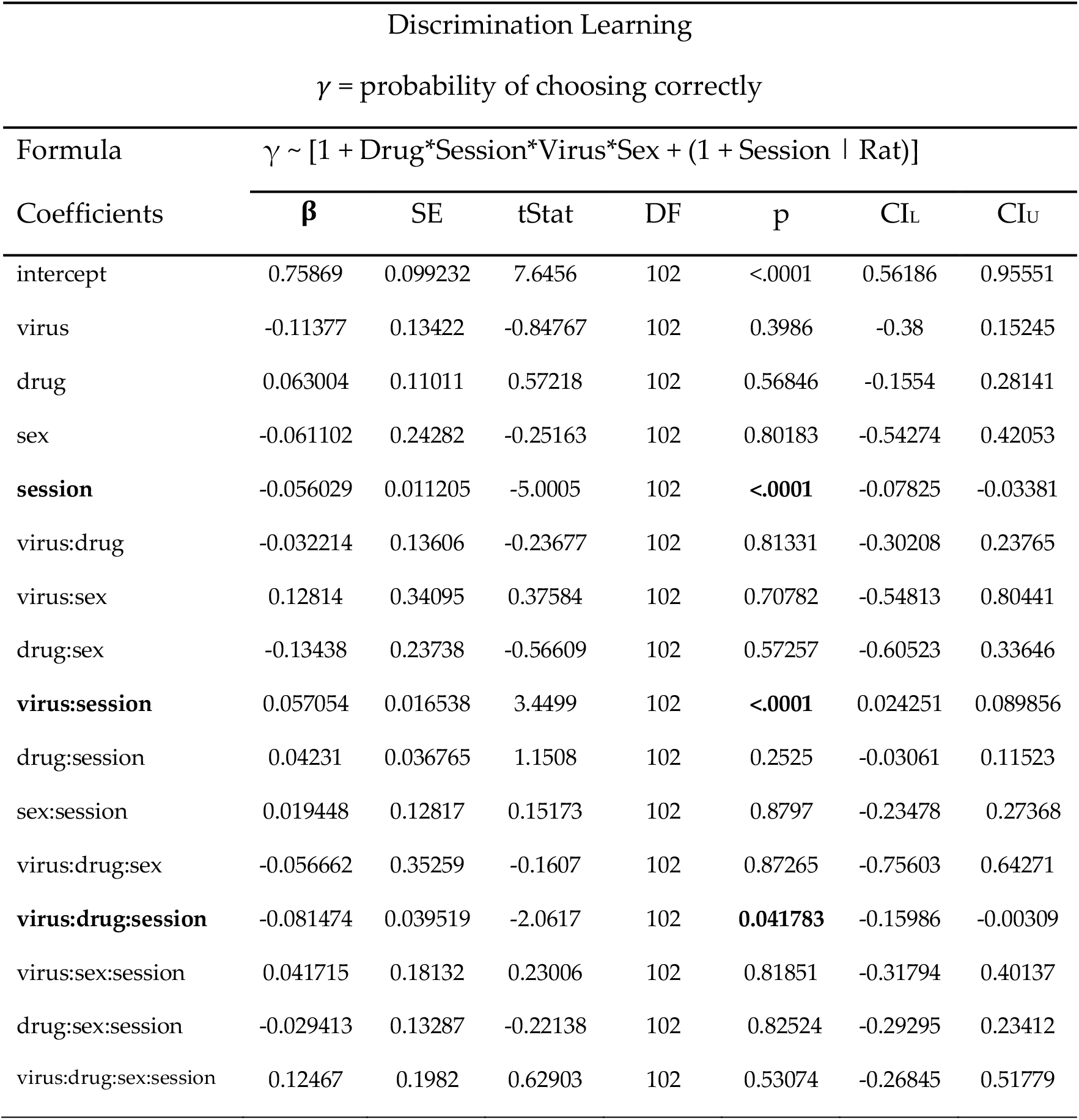
Probability of choosing correctly during initial visual discrimination learning.

**Table 2.**
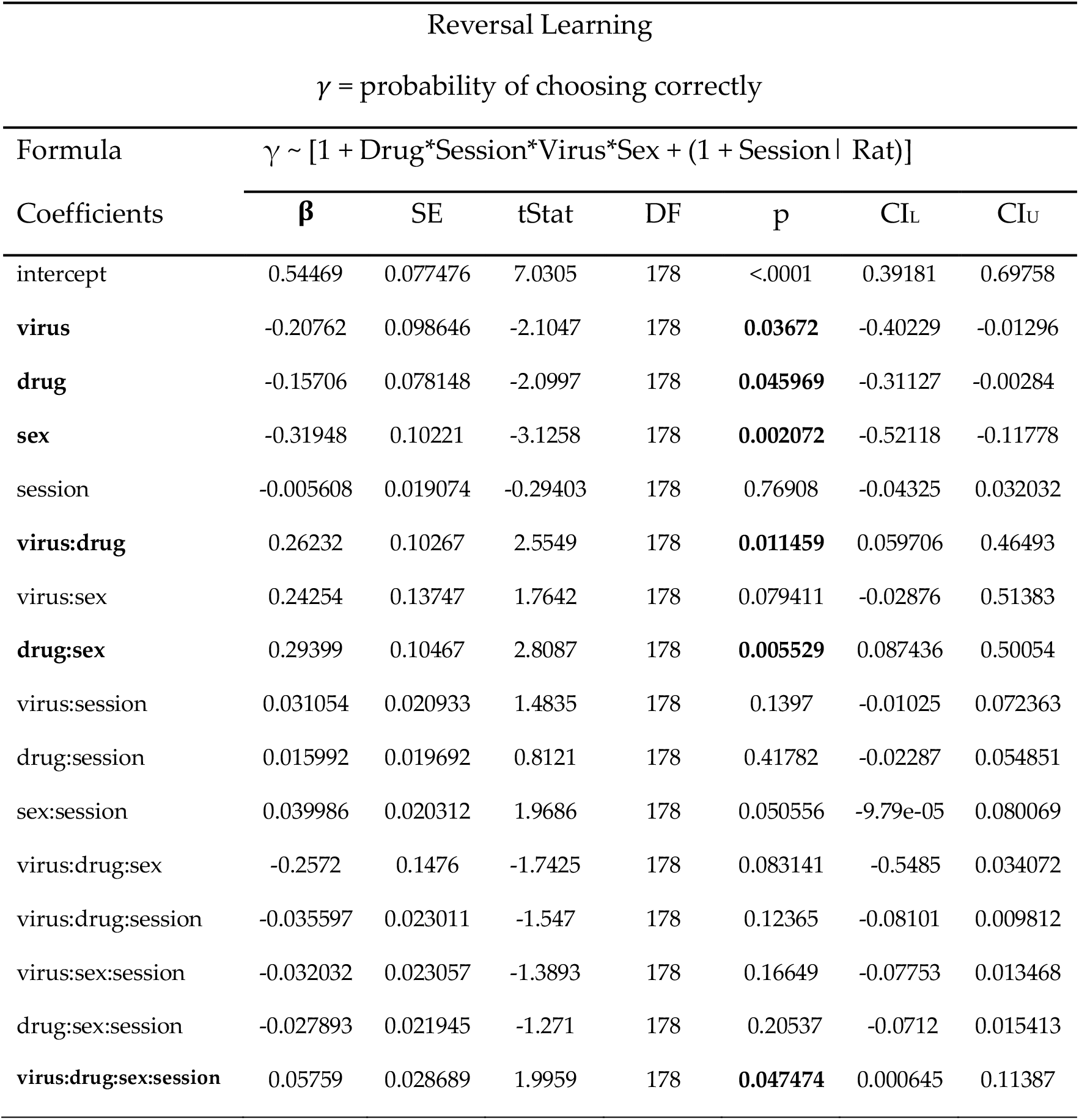
Probability of choosing correctly during reversal learning.

**Figure 2.**
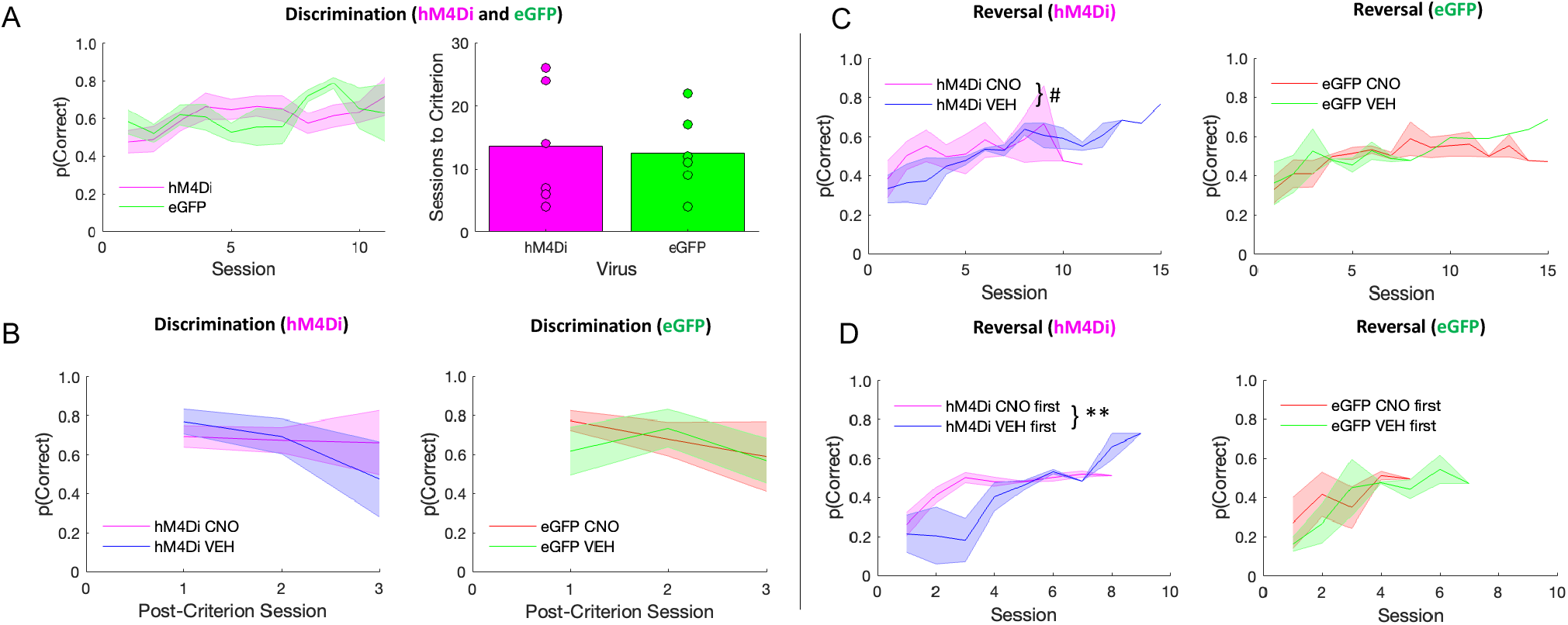
ACC inhibition has no effect on discrimination performance, but promotes random responding during early reversal learning. There were no differences between hM4Di and eGFP virus groups in the initial discrimination phase, however, clozapine-N-oxide (CNO) affected long-term reversal learning differently depending on virus group, and depending on when CNO was administered. **(A)** Performance in hM4Di rats and eGFP rats during initial discrimination of stimuli during which there was no recording and no drug administered (*left*). Rats did not differ by virus group in the number of sessions to reach criterion (*right*). **(B)** There was no difference between virus groups or by drug in post-criterion discrimination performance. **(C)** Learning curves during the reversal phase split by hM4Di (*left*) and eGFP (*right*). irrespective of virus condition, rats rarely surpassed 60% accuracy. **(D)** Rats received either VEH or CNO first during the reversal phase prior to reaching 50% correct. Same learning as in C, but learning curves now split by whether rats received CNO first or VEH first. ACC inhibition during early reversal learning promotes a random response strategy that they do not recover from. Means ± standard error of the mean (S.E.M.) shades. Post-hoc Bonferroni comparisons, #p=0.07, **p=0.001. hM4Di n=8 (4 female), and eGFP n=7 (3 female).

### ACC inhibition attenuates a trial accuracy theta signal in OFC during discrimination

In a subset of learners, we collected local field potential (LFP) data in ACC and OFC (**Figure 3**), acquired at 30 kHz and down sampled to 1,000 Hz, and bandpass filtered for theta (5-10 Hz). We were primarily interested in measures of accuracy: correct vs. incorrect trials (Harris, Aguirre et al. (2021), but we also included trial initiation and reward collection latencies in our analyses. The latter measures did not emerge as significant correlates or predictors of learning.

**Figure 3.**
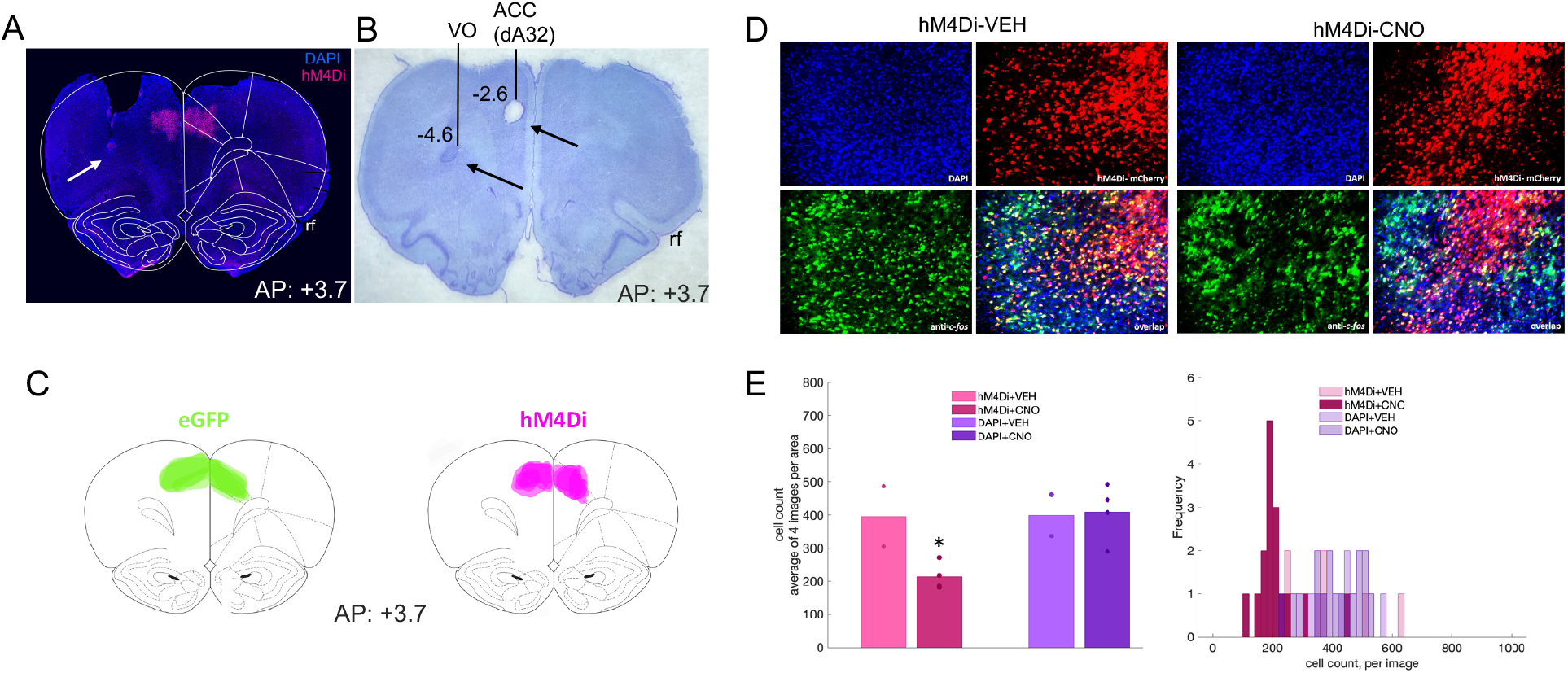
Inhibitory DREADDs in ACC, electrode placement in ACC and OFC, and validation of inhibition in ACC via c-fos immunohistochemistry. **(A)** Representative placement of inhibitory hM4Di DREADDs at Anterior-Posterior (AP) level +3.7 relative to Bregma. Note, tissue (secondary motor cortex, M2) was removed at time of brain extraction. White arrow pointing at tip of electrode track in OFC. **(B)** Nissl-stained section showing electrolytic lesions and placement of electrode arrays targeting both ACC (dorsal area 32, dA32) and OFC (ventral orbital, VO) unilaterally in a representative animal. Black arrows pointing at tip of electrode tracks. rf = rhinal fissure. **(C)** Reconstructions of placement of inhibitory hM4Di DREADDs (right) and eGFP null virus (left) at Anterior-Posterior (AP) level +3.7 relative to Bregma. **(D)** Representative DAPI, hM4Di-mCherry, c-fos immunoreactivity and their overlap after injections of VEH (left) or CNO (right). **(E)** Mean cell count of 4 images per condition, hM4Di+VEH, hM4Di+CNO, DAPI+VEH, and DAPI+CNO. One-way ANOVA resulted in a significant decrease in c-fos positive cells in the hM4Di-expressing areas, but not in the DAPI areas (left). Histogram of cell count frequency for each image by condition (right). *p<0.05.

Based on the temporal profile of theta power changes during behavior in animals transfected with hM4Di (**Figure S2)** and eGFP in ACC (**Figure S3**), we baseline-subtracted theta band in ACC and OFC using the immediate pre-event period (−200 ms) leading up to the choice of the correct or incorrect stimulus. This was the moment at which the animal had already initiated a trial, and was fixating on the stimulus of choice. We calculated this for both discrimination and reversal phases. Despite no performance decrements following ACC inhibition in discrimination learning, normalized theta power in ACC during both correct and incorrect trials was significantly lower after CNO than after VEH in hM4Di animals (**Figure 4A**). We performed 2 × 2 ANOVAs on this baseline-subtracted theta power comparing trial type (correct, incorrect) and drug (VEH, CNO) for discrimination sessions, separately for OFC and ACC. In ACC, we observed a significant effect of trial type [*F*(1,45)=14.237, *p*<0.0001] and a significant drug x trial type interaction [*F*(1,45)=5.859, *p*=0.02], with increases in ACC theta in both correct and incorrect trials in the VEH compared to the CNO condition (Bonferroni-corrected post-hoc comparisons, all *p*<0.035). There was no such pattern for normalized OFC theta during discrimination, and there was a nonsignificant effect of drug [*F*(1,50)=3.965, *p*=0.052] (**Figure 4B**). In animals with eGFP virus, one could distinguish correct from incorrect trials from normalized theta in OFC [trial type x drug: *F*(1,27)=6.940, *p*=0.01, Bonferroni-corrected post-hoc comparisons, all p<0.01], but not in ACC. In sum, ACC inhibition attenuated a trial accuracy theta signal in OFC during discrimination (**Figure 4B** when compared to **Figure 4D**), but these changes did not appreciably impact accurate performance.

**Figure 4.**
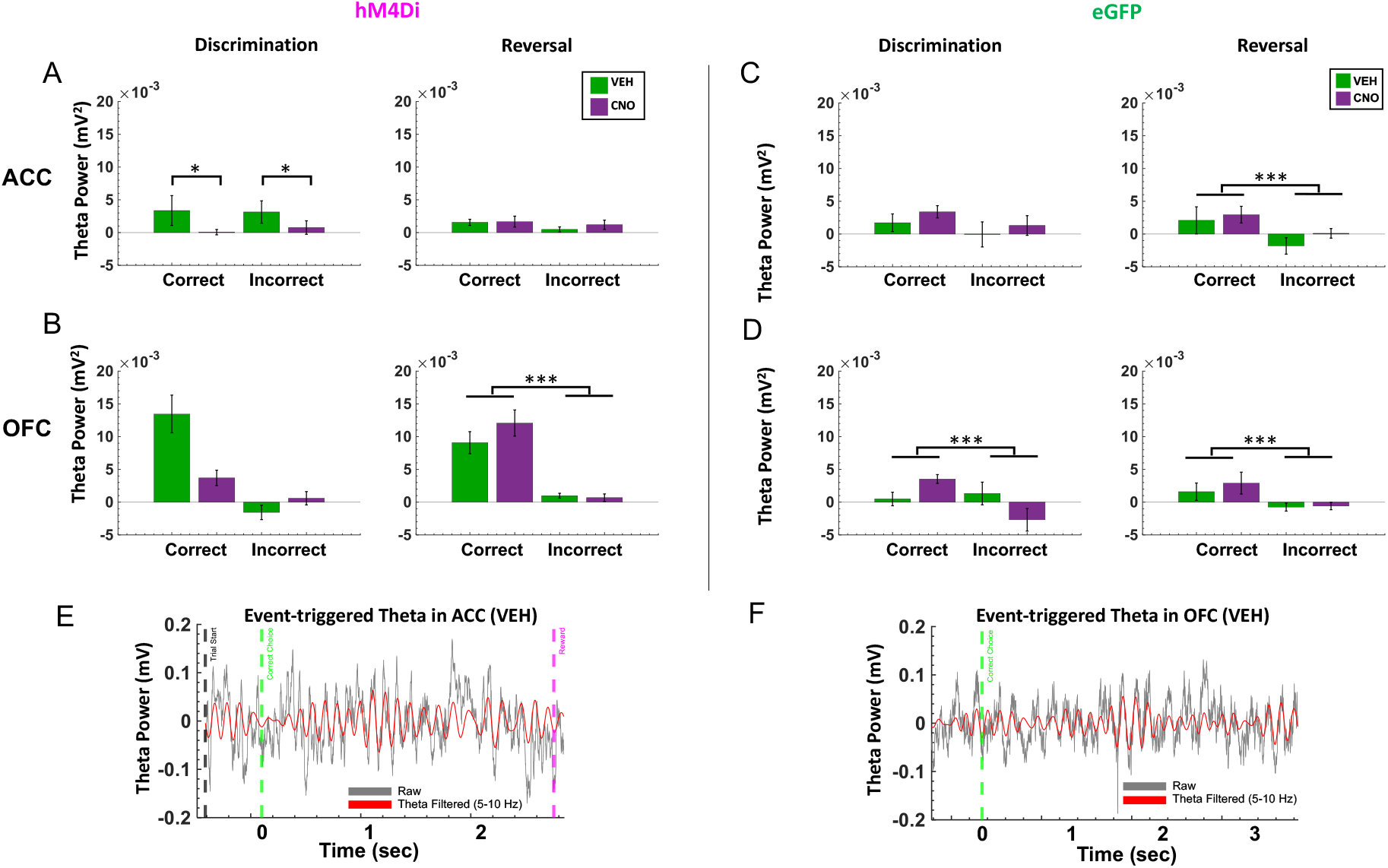
Trial accuracy can be distinguished only from OFC, not ACC, theta power in the reversal phase following ACC inhibition. Baseline-subtracted (normalized) theta power was calculated for correct and incorrect choices in the discrimination and reversal learning phases. (**A**) In hM4Di animals, there was no significant difference in normalized ACC theta power between correct vs. incorrect trials in the reversal phase. (**B**) In hM4Di animals, there was no significant difference in normalized theta power in OFC for correct vs. incorrect trials in discrimination, but normalized theta power in OFC did signal trial accuracy in reversal learning. (**C)** In eGFP animals, normalized theta power in ACC signaled trial accuracy in reversal learning. (**D**) In eGFP animals, normalized theta power in OFC signaled trial accuracy in both discrimination and reversal learning. (**E**) Example event-triggered raw (gray) and theta filtered (red) trace in hM4Di rat with trial start (black dotted line) and correct choice (green dotted line), and reward collection (pink dotted line) denoted for the trial. (**F**) Example event-triggered raw (gray) and theta-filtered (red) trace in eGFP rat with correct choice (green dotted line) denoted for the trial. Means ± standard error of the mean (S.E.M.) bars. ***p<0.001, *p<0.05 following Bonferroni-corrected post-hoc comparisons.

### Correct vs. incorrect trials are strongly differentiated by OFC, but not ACC, theta power in the reversal phase following ACC inhibition

Following ACC inhibition, correct vs. incorrect trials in reversal learning could only be distinguished by OFC, not ACC, normalized theta. As above, we performed two separate 2 × 2 ANOVAs, comparing trial type (correct, incorrect) and drug (VEH, CNO) for all reversal sessions, separately for OFC and ACC. In hM4Di animals, there was a main effect of trial type on normalized OFC theta in reversal [*F*(1,143)=51.250, *p*<0.0001, all corrected Bonferroni post-hoc comparisons of correct vs. incorrect trials, all *p*<0.001] (**Figure 4B**). This difference by trial type in normalized theta power was absent in ACC, in the hM4Di animals. In eGFP animals, there was a significant effect of trial type for normalized OFC theta [*F*(1,132)=6.21, *p*<0.01] and ACC theta [*F*(1,132)=6.109, *p*=0.01], with post-hoc Bonferroni-corrected comparisons revealing normalized theta was greater for correct choices compared to incorrect choices (all post-hoc tests, *p*<0.001), **Figure 4CD**. Thus, ACC inhibition reduced a trial accuracy theta signal during reversal learning in ACC (**Figure 4A** compared to **Figure 4C**), but had no impact on OFC theta. Results plotted by early vs. late reversal revealed a similar pattern (**Figure S4**): there was a main effect of trial type on normalized OFC theta in reversal, whether rats received VEH first [*F*(1,52)=37.235, *p*=1.33e^-07^] or CNO first [*F*(1,70)=17.88, *p*=6.99e^-05^]; all Bonferroni corrected post-hoc of correct vs. incorrect trials, *p*<0.001. Though there was the same trend for OFC theta in eGFP animals, this difference was not statistically significant. Additionally, an analysis of individual rat (not session) changes in normalized theta revealed the same pattern of an attenuated trial accuracy theta signal during reversal learning in ACC, and an intact OFC theta signal of correct vs. incorrect in the hM4Di group (**Figure S5**): [*F*(1,15)=20.831, *p*=6.50e^-04^]; all Bonferroni corrected post-hoc of correct vs. incorrect trials, *p*<0.001.

### ACC inhibition hastens incorrect choices in reversal learning

Because we were also interested in theta oscillations as they relate to speed of response, we next analyzed median latencies, i.e., to initiate a trial, to choose the correct stimulus, to choose the incorrect stimulus, and to collect reward (Harris, Aguirre et al. 2021) in both discrimination and reversal learning. We focused on measures for which an omnibus across-phase mixed GLM analyses revealed significant phase and drug interactions: *incorrect choice latencies* and *reward collection latencies*.

In hM4Di animals for the discrimination phase, drug, ACC theta, and OFC theta were not significant predictors of incorrect choice latencies (GLM formula: *γ* ∼ [*1 + Drug*\**ACCtheta* \**OFCtheta + (1+ Drug)*]). However, for the reversal phase, we found a significant effect of drug (β_drug_ =-4.062, t(67)=-2.289, *p*=0.025; VEH slower than CNO, respective means 1.88s vs 1.72s, **Figure 5A**), and OFC theta, not ACC theta, was a significant predictor of incorrect choice latencies (GLM: β_OFCtheta_ =-89.086, t(67)=-2.377, *p*=0.020). For reward collection latencies in the reversal phase, we found a significant effect of drug (GLM: β_drug_ =-4.217, t(67)=-2.081, *p*=0.041; VEH slower than CNO, respective means 0.63s vs 0.52s). We also observed a significant negative correlation between OFC theta power and reward collection speed under CNO (r=-0.14934, *p*=0.031) but not under VEH conditions (r=-0.0071, *p*=0.911). There were no other correlations between theta power and speed in hM4Di rats (**Figure 6**).

**Figure 5.**
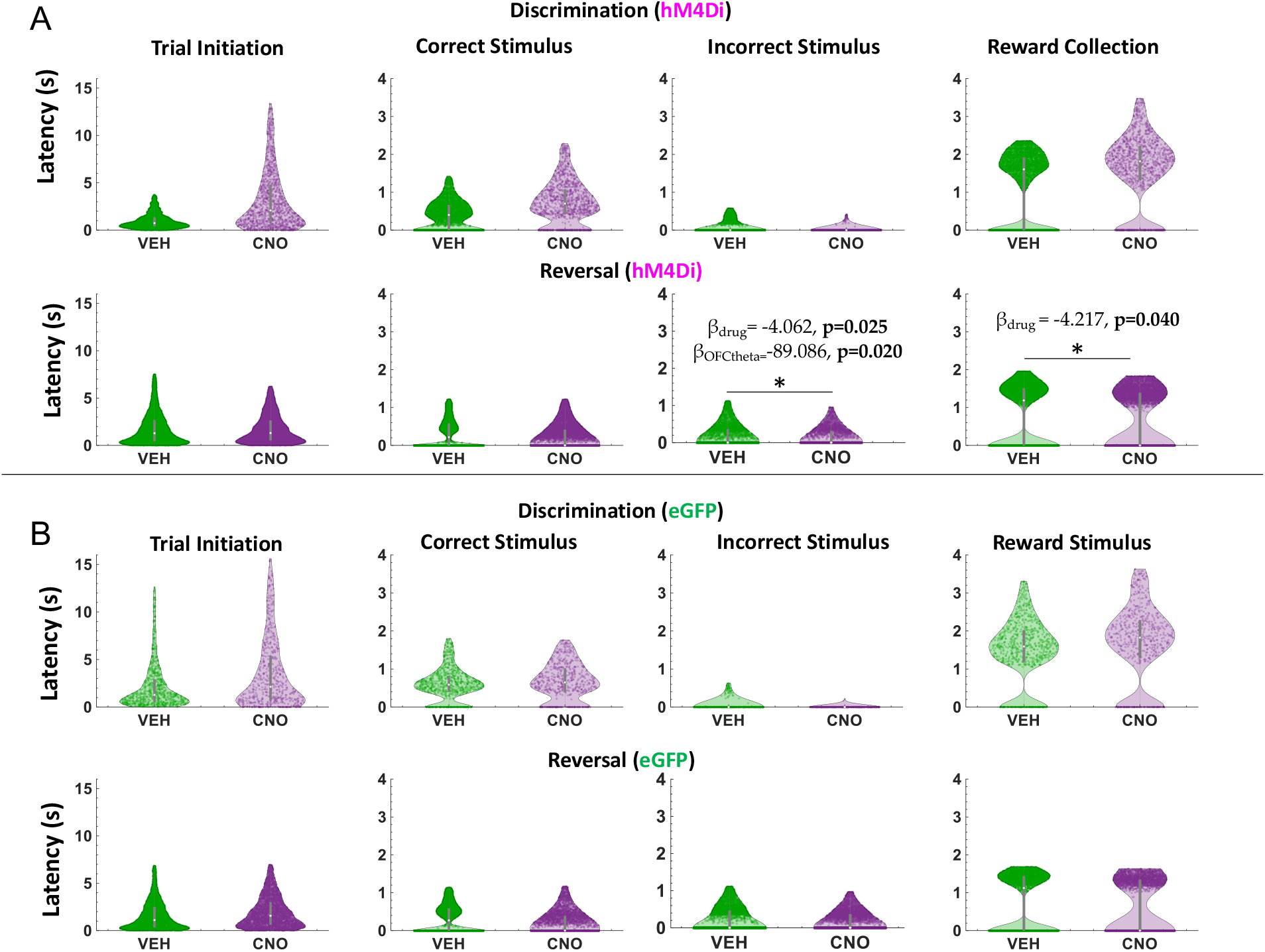
ACC inhibition increases speed of incorrect choices in reversal learning. Violin plots depicting individual trial median latencies in seconds (s) for each of the trial epochs in discrimination and reversal learning for hM4Di and eGFP rats. Individual dots represent trials. (**A**) In hM4Di rats, clozapine-N-oxide (CNO) administration significantly reduced the latency to choose the incorrect stimulus and collect reward in the reversal, not discrimination, phase. ACC theta did not predict speed of responses for any trial epoch in either the discrimination or reversal phase. (**B**) In eGFP rats, there was no significant hastening of incorrect choices, though there was a significant drug x phase interaction for trial initiation latencies (subsequent post-hoc effects were not statistically significant). All significant Beta values are not shown, but are reported in text. *p<0.05, CNO vs. VEH.

**Figure 6.**
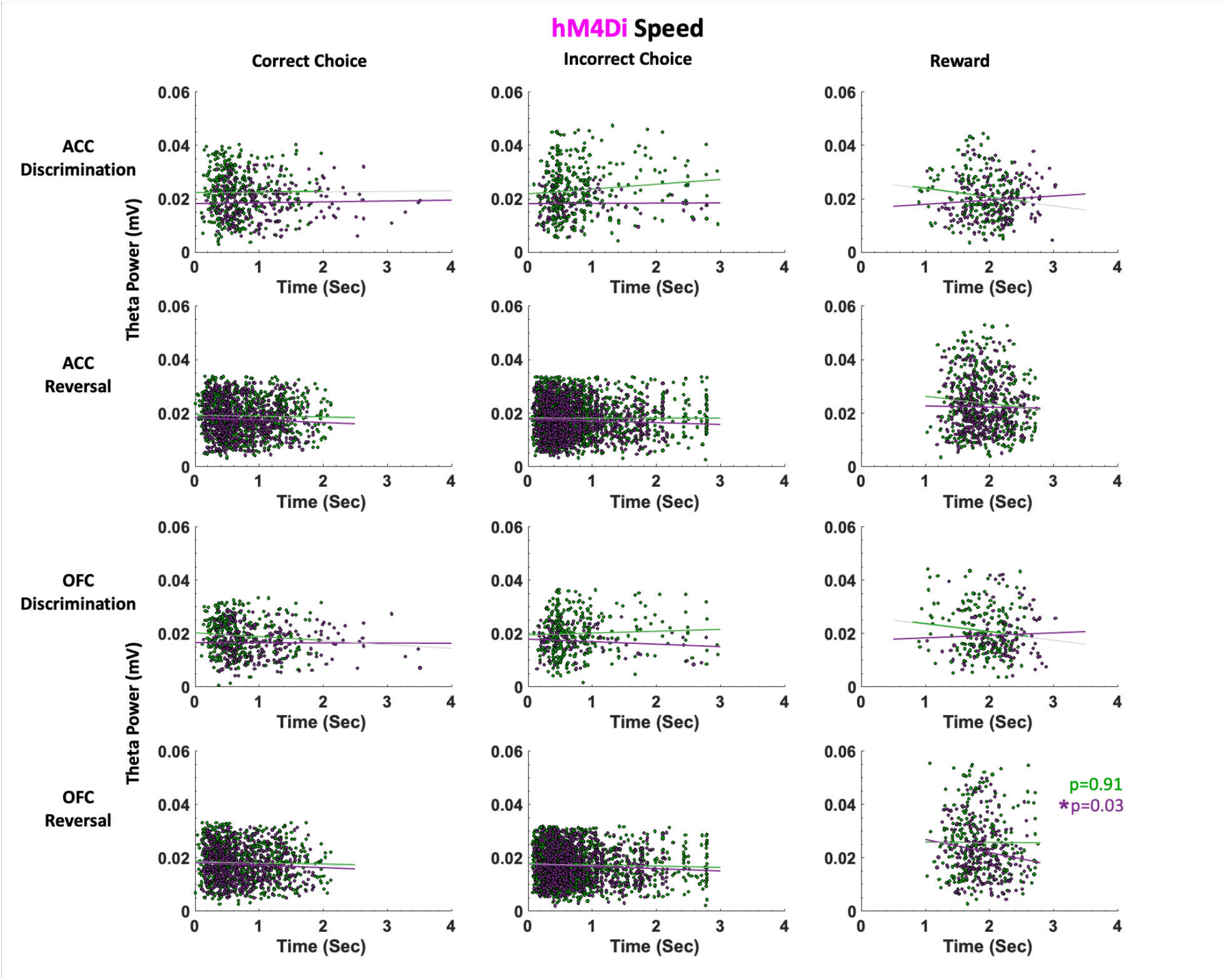
ACC and OFC theta power are not correlated with deliberation speed in either discrimination or reversal learning in hM4Di rats. Theta power was filtered and extracted from ACC and OFC for each event-triggered epoch (Correct choice, Incorrect choice, and Reward port entry). Movement speed was calculated by subtracting the time at the target event (t) from the time at the previous event (t-1) in seconds (Sec). OFC theta power decreased as reward collection latencies increased after CNO administration (purple) during reversal learning. This negative correlation was not observed following VEH (green). There were no significant correlations between theta power and choice of correct or incorrect stimulus. *p<0.05.

In eGFP rats, we did not observe significant differences in median latencies when comparing CNO vs. VEH conditions in either the discrimination or reversal phase (**Figure 5B**). Also unlike in hM4Di animals, in eGFP rats there were significant correlations between speed of correct choices (i.e., deliberation speed) and theta power in the VEH, but not CNO, conditions. During post-criterion discrimination performance, OFC theta power was negatively correlated with correct choice (r=-0.13, p=0.02), but not under CNO conditions (r=-0.05, p=0.331). Similarly, ACC theta power was negatively correlated with deliberation speed for correct choices under VEH (r=-0.11, p=0.049), but not under CNO (r=-0.03, p=0.56). In reversal learning, both ACC theta power (r=-0.16, *p*=3.55e-08) and OFC theta power (r=-0.08, *p*=0.01) were negatively correlated with reward collection speed under CNO, but not VEH, conditions (**Figure 7**). We also found a significant drug x phase interaction (GLM: β_drug X phase_ =-3.329, t(65)=-2.301, *p*=0.025) for initiation latencies, however post-hoc analyses did not reveal any statistically significant differences.

**Figure 7.**
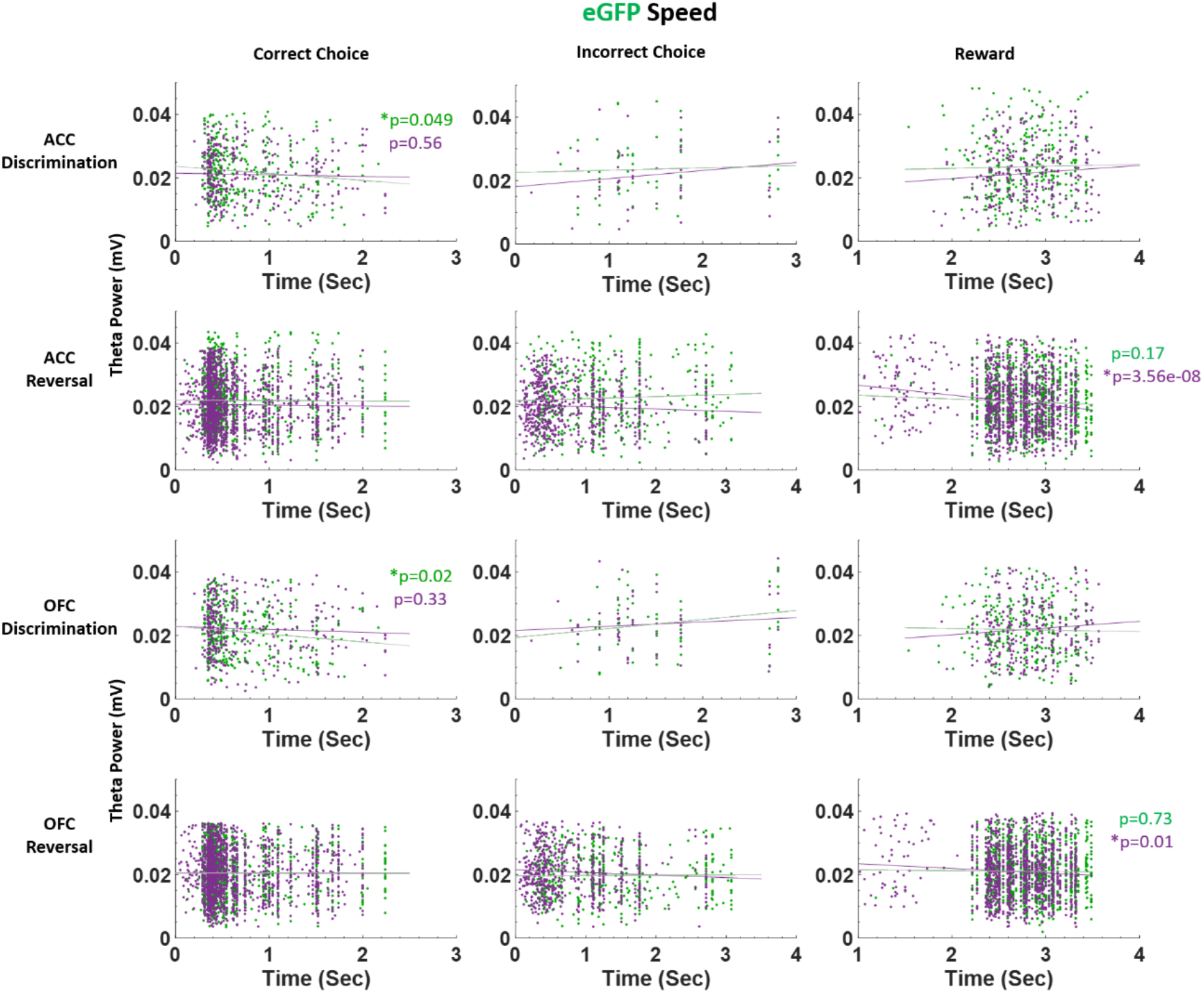
ACC and OFC theta power are negatively correlated with deliberation speed during discrimination performance in eGFP rats. Theta power was filtered and extracted from ACC and OFC for each event-triggered epoch (Correct choice, Incorrect choice, and Reward port entry). Movement speed was calculated by subtracting the time at the target event (t) from the time at the previous event (t-1) in seconds (Sec). OFC theta power decreased as correct choice latencies increased after VEH administration (green) during discrimination learning. This negative correlation was not observed following CNO (purple). In reversal learning, theta power in both ACC and OFC significantly decreased as reward collection latencies increased after CNO administration (purple). This negative correlation was not observed following VEH (green). There were no significant correlations between theta power and choice of correct or incorrect stimulus in reversal learning. *p<0.05.

Our finding speed-theta power correlations for correct choices in eGFP, and not hM4Di, animals suggest control rats are more certain of correct choices (i.e., faster correct choice, greater theta power) under VEH than under CNO during post-criterion performance, and further, that CNO enhances the relationship between reward collection speed and theta power in OFC of control rats.

Taken together, the latency analyses in learning reveal quicker deliberation speed for incorrect trials during the reversal phase following ACC inhibition. Notably, CNO had an effect on reward collection times in both eGFP and hM4Di animals, with greater theta power in OFC associated with quicker reward collection times, indicating a drug effect, not an ACC inhibition effect. However, reward collection latencies best indicate motivation levels, not deliberation speed (Harris, Aguirre et al. 2021). We show here that CNO does not produce global locomotor effects which would be reflected across all trial epochs in our task, in contrast to a previous report (MacLaren, Browne et al. 2016).

### Theta coherence during correct vs. incorrect trials

Previous studies by other groups have explored the degree to which the ACC and OFC are synchronized during effort- and value-based decisions (Fatahi, Haghparast et al. 2018, Fatahi, Ghorbani et al. 2020), with evidence suggesting significant coherence in the theta-band prior to, and after, making the correct decision. Given that we observed differences in choice accuracy between hM4Di and eGFP animals, coherence analyses during these triggered events could provide a deeper understanding of ACC-OFC theta synchronization. Here, spectral coherence was calculated across trials after aligning the raw local field data from −2 to +2 seconds of the animal’s choice, with time=0 indicating the precise time in which the choice was made. Identical to the theta-band filtered data above, the average pre-choice period (−200 to 0 ms) was subtracted from the entire −2 to +2 second window. In the discrimination phase, a 2×2 (drug x virus) ANOVA revealed a significant difference between hM4Di vs. eGFP virus groups in theta coherence during correct choices in the discrimination phase [*F*(1,39)=6.538, *p*=0.01]. However, Bonferroni-corrected post hoc *t*-tests did not result in a significant difference. We did not observe any statistically significant coherence differences for incorrect choices during discrimination phase (**Figure S5**) or for correct or incorrect choices in the reversal phase (**Figure S6**). Thus, we found ACC-OFC theta coherence to be fairly weak in our task despite observing significant differences in theta power during choice accuracy (**Figure 4**).

## DISCUSSION

In the present study we were specifically interested in theta oscillations in frontal cortex since there have been several recent reports of involvement of oscillations in this frequency band in reward value and decision making in rodents (Marquardt, Sigdel et al. 2017, Fatahi, Haghparast et al. 2018, Fatahi, Ghorbani et al. 2020, Amarante and Laubach 2021), and one closed-loop experiment in non-human primates demonstrating that theta oscillations in OFC are causally involved in reversal learning (Knudsen and Wallis 2020). We found strong support for OFC theta signaling of accuracy in reversal learning. Theta is a large-scale network oscillation believed to synchronize interregional communication (Buzsaki 2005). The precise origins of OFC theta oscillations are unknown but theta oscillations propagating from the hippocampus to the OFC have been found to encode value learning in macaques (Knudsen and Wallis 2020). While hippocampal theta in rodents is well-established to be associated with certain aspects of movement (Kennedy, Zhou et al. 2022), others have shown that the strength of theta-band phase locking in OFC neurons follows the rat’s current outcome expectation, which can be dissociated from licking responses (van Wingerden, Vinck et al. 2010). While volume conduction from the hippocampus is a possibility, the placement of our electrodes in two areas of prefrontal cortex supports a cortical origin for the measured theta oscillations. Previous findings of OFC theta phase locking of spiking patterns also argue against simple hippocampal volume conduction (van Wingerden, Vinck et al. 2010), as well as evidence that ACC coherence with hippocampus is dynamically modulated by behavior, such that ACC theta is not just an attenuated copy of hippocampal theta (Young and McNaughton 2009, Remondes and Wilson 2013).

Here we observed a robust OFC theta power signal of accuracy in reversal learning sessions in eGFP and hM4Di animals, unperturbed by ACC inhibition. Since LFPs in OFC, but also broadly in other areas, constitute a composite signal of multiple afferents, it is likely that another input other than ACC is more crucial to the involvement of OFC theta in reversal learning. Additionally, establishing a causal role for oscillations using chemogenetic or other viral approaches is difficult because such manipulations likely disrupt underlying firing rates, a limitation that could be bypassed by a closed-loop approach (Knudsen and Wallis 2020). In particular, the specific silencing of CaMKII projection neurons is not a natural physiological state, and this may trigger compensatory mechanisms in downstream targets that should be investigated in high-yield recording studies in the future.

We also analyzed ACC-OFC coherence and found no evidence of significantly enhanced coherence during correct vs. incorrect choices in our task. Other groups have previously utilizeda t-maze task that incorporates a spatial component to the decision process, with necessary running both prior to as well as after the choice point in the maze (Fatahi, Haghparast et al. 2018, Fatahi, Ghorbani et al. 2020). While these authors did not determine whether running speed was correlated with ACC-OFC coherence on correct trials, it remains a possibility. High theta coherence between ACC and hippocampus has been associated with coding of an animal’s current position during trajectories along a maze that may contribute to decision making (Remondes and Wilson 2013, Remondes and Wilson 2014, Zielinski, Shin et al. 2019). Unlike these maze tasks, movement was much more limited in our task. Indeed, we observed the largest changes in theta power when animals made the choice of the correct stimulus, at which time they were most stationary, just having initiated a trial while fixating on the stimulus. Additionally, we found there were significant increases in speed of incorrect choices following ACC inhibition in the same (reversal) phase when there was no theta ‘read-out’ of correct vs. incorrect trials. This provides some evidence that ACC inhibition eroded the distinction between correct and incorrect stimuli in the reversal phase, further elaborated below.

Various studies, mostly rodent lesion and pharmacological inactivation studies, support the idea that ACC is involved in initial cue or stimulus discrimination learning (Chudasama, Passetti et al. 2003), but has little-to-no involvement in fully-predictive, deterministic reversal learning (Bussey, Muir et al. 1997, Schweimer and Hauber 2006). Conversely, the prediction based on the literature is the opposite for OFC: critical for reversal, but not initial discrimination learning (Schoenbaum, Chiba et al. 1999, Marquardt, Sigdel et al. 2017). Thus, the question about if and how ACC may be causally-involved in stimulus-based reversal learning was a question ripe for investigation. To our knowledge, we are the first to probe this using visual stimuli and with a viral-mediated approach targeting principal neurons in rats. We found that chemogenetic inhibition of ACC indeed impacts reversal learning. The quick adoption of 50% responding is interesting and suggests that animals are either 1) newly satisfied with a 50% reward rate, or 2) unable to distinguish a discrimination vs. reversal ‘task state’, and thus adopt a random response strategy. The latter possibility is more likely since we found no consistent evidence of motivational changes following ACC inhibition. Labeling of the current task state and building of a cognitive map has been linked to OFC (Wilson, Takahashi et al. 2014, Costa, Scholz et al. 2022), yet our data suggest it may also be a role of ACC, at least in rats. The latency data we report here (i.e., demonstrating that animals do not deliberate differently between correct and incorrect stimuli), support this possibility. Surprisingly, we found a drug x virus interaction in reversal learning such that CNO had a dissociable, but non-negligible effect in eGFP reversal learning. Given that this effect of CNO was specific to the late reversal phase and not observed in discrimination learning, it suggests that administration of CNO when performance is at chance may further disrupt reversal learning, given that reversal learning is modulated via a dopamine D2 mechanism (Baldessarini, Centorrino et al. 1993, Lee, Groman et al. 2007, Linden, James et al. 2018, Manvich, Webster et al. 2018).

As suggested above, theta oscillations in the hippocampus have been shown to correlate with cognitive processing, movement speed and acceleration (Vanderwolf 1969, O’Keefe and Nadel 1978, Kennedy, Zhou et al. 2022). Some evidence suggests theta oscillations in frontal cortex promote inter-regional coordination (Jones and Wilson 2005, Siapas, Lubenov et al. 2005, Cavanagh and Frank 2014, Rajan, Siegel et al. 2019), yet the nature of these relationships is much less firmly established across subregions of frontal cortex. Because our rats are freely-behaving in the operant chambers, we similarly assessed speed of responding at specific trial epochs and correlations with theta power in ACC and OFC. We found that ACC inhibition hastened incorrect choices in reversal learning: this was observed only in hM4Di animals and not in eGFP animals, indicating it was indeed a result of ACC inhibition and not a broad drug effect. In a mixed-effects GLM model with ACC theta, OFC theta, and drug as predictors of latencies, OFC theta and drug (CNO) emerged as the only significant predictors of these quick incorrect choices. In follow-up work, other frequency bands could be systematically probed for speed correlations since not only theta (Fatahi, Haghparast et al. 2018, Fatahi, Ghorbani et al. 2020) but also gamma (Donnelly, Holtzman et al. 2014) oscillations in frontocortical regions have been linked to preparatory motor responses to cues that predict reward. Similarly, beta increases (Schmidt, Herrojo Ruiz et al. 2019) have also been observed at trial-end when working memory engagement is high (in the absence of discrete cues).

In summary, the present results suggest a role for ACC in stimulus-based reversal learning; a role similar to that historically proposed for OFC. As several psychiatric conditions manifest impairments in reversal learning including substance use disorder, obsessive compulsive disorder, and schizophrenia (Remijnse, Nielen et al. 2006, Brigman, Ihne et al. 2009, Leeson, Robbins et al. 2009, Winstanley, Olausson et al. 2010, Izquierdo and Jentsch 2012), frontal cortex theta oscillations could be studied as biomarkers in preclinical models of these disorders.

## MATERIALS AND METHODS

### Animals

A total of N=24 Long-Evans rats (Charles River Laboratories, Hollister, CA) was used for these experiments. Three animals were not included in the final dataset due to implant failure. Six male animals were surgerized to express hM4Di and after three weeks were euthanized shortly after being administered 3 mg/kg of clozapine-N-oxide (CNO), to assess the number of c-fos positive cells in hM4Di-expressing areas versus neighboring (DAPI) areas following vehicle and CNO administration (see details below). Of the n=15 rats used for learning, eight were used for electrophysiological recordings. The latter includes male (*n*=4) and female (*n*=4) animals. All rats were post-natal day 40 (∼250g) upon arrival and were pair-housed in a 12-h reverse light/dark cycle room (lights on at 06:00h), maintained in 22-24ºC temperature conditions, with food and water available *ad libitum* prior to behavioral testing. After one week of acclimation to our vivarium, all animals were handled for 10 min in pairs for 5 days. After the handling period, animals were individually-housed to carefully monitor food consumption under restricted access. All procedures shown in the experimental timeline (**Figure 1A**) were in accordance with the recommendations of the Guide for the Care and Use of Laboratory Animals of the National Institutes of Health. The protocol was approved by the Chancellor’s Animal Research Committee at the University of California, Los Angeles.

### Viral Constructs

An adeno-associated virus AAV8 driving the hM4Di-mCherry sequence under the CaMKIIa promoter was used to express DREADDs on putative projection neurons in ACC (*AAV8-CaMKIIa-hM4D(Gi)-mCherry*, packaged by Addgene, Addgene, viral prep #50477-AAV8). A virus lacking the hM4Di DREADD gene and instead containing the fluorescent tag eGFP (*AAV8-CaMKIIa-EGFP*, packaged by Addgene) was infused into ACC in 4 animals as a null virus control. The DREADDs-transfected and null eGFP animals underwent identical surgeries, and all of our behavior was conducted in a counterbalanced within-subject design with each animal serving as its own control (VEH compared to CNO). Collectively, this allowed us to control for non-specific effects of surgical procedures, exposure to AAV8, and non-specific effects of CNO. We note that several recent electrophysiology experiments combining neural recordings with DREADDs do not include virus (eGFP) controls (Liu, McAfee et al. 2017, Alexander, Brown et al. 2018, Schmidt, Duin et al. 2019), as we do here.

### Behavioral Apparatus

Behavioral testing was conducted in operant conditioning chambers shielded for electrophysiological recordings and outfitted with an LCD touchscreen opposing a sucrose pellet dispenser (Lafayette Instrument Co., Lafayette, IN). Sucrose pellets (45mg, Dustless Precision Pellets #F0023, Bio-Serv) were used as rewards. The inner chamber walls were modified to accommodate implanted animals by maximizing the space available (24cm × 33.2cm × 39.5cm) and to improve electrophysiological signal quality. All chamber equipment was controlled by customized ABET II TOUCH software.

### Pretraining

Immediately following the 5 days of handling, rats were placed on food restriction (12 g/d) to no less than 85% of their free-feeding body weight throughout the entire experiment and were weighed each day and monitored closely to not fall below this percentage. Animals were pretrained using the protocol as previously published (Stolyarova and Izquierdo 2017). Briefly, a series of pretraining stages: Habituation, Initiation Touch to Center (ITC), and Immediate Reward (IM), were designed to train rats to nosepoke, initiate a trial, and select a stimulus to obtain a reward.

In the habituation stage, five pellets were automatically dispensed and rats were required to consume all pellets within 15 min to advance. In the ITC stage, the center of the touch screen displayed a white square graphic against a black background. A single sucrose pellet was dispensed with simultaneous onset of audio tone and illumination of the reward receptacle if the rat nosepoked the white stimulus, if rats took longer than 40 s to take this action, the white stimulus disappears without reward and trial was considered an omission followed by a 10 s inter-trial interval (ITI). Rats were required to reach a criterion of consuming 60 rewards in 45 min in order to advance to IM training. In this stage, the same white stimulus was presented as in ITC and a nosepoke indicated trial initiation. The disappearance of the white stimulus was immediately followed by the presentation of a target stimulus on the left or right side of the touchscreen (i.e., forced choice), nose poking this target stimulus was paired with a reward. An *initiation omission* was scored if rats took longer than 40 s to initiate the trial, a *choice omission* was scored if rats failed to nosepoke the left or right stimulus after 60 s. A criterion of 60 rewards consumed in 45 min across two consecutive days was required to advance to the discrimination phase of the behavioral task.

### Surgical Procedures

After completing the final pretraining stage, rats were anesthetized with isoflurane for bilateral ACC DREADDS (AAV8-CaMKIIα-hM4D(G_i_)-mCherry, Addgene, Cambridge, MA, Addgene, viral prep #50477-AAV8) or eGFP (AAV8-CaMKIIa-EGFP, Addgene, Cambridge, MA, Addgene, viral prep #50469-AAV8) infusion and unilateral chronic implantation of electrode arrays in both ACC and OFC. Craniotomies were created and a 26-gauge guide cannula (PlasticsOne, Roanoke, VA) were lowered into ACC (AP: +3.7, ML: ±0.8, DV: −2.6 from skull surface), after which a 33-gauge internal cannula (PlasticsOne, Roanoke, VA) was inserted. This set of coordinates for ACC was the more anterior site of the two sites that our lab has previously utilized: more posterior ACC has been probed in effort-based tasks (Hart, Blair et al. 2020) whereas stimulus-based reversal learning has been probed with this more anterior targeting (Stolyarova, Rakhshan et al. 2019), constituting area 32 of cingulate cortex (van Heukelum, Mars et al. 2020). Animals were infused with 0.3 μL of virus at a flow rate of 0.1 μL/minute, with cannulae left in place for 5 additional minutes to allow for diffusion.

In the same surgery after DREADDs infusion, rats were implanted with a custom-designed 16-channel electrode array manufactured in-house. The array was composed of 8 stereotrodes (California Fine Wire Co., Grover Beach, CA) with four targeting the ACC and another four aimed simultaneously in the ipsilateral OFC (AP: +3.7, ML: ±2.0, DV: −4.6, from skull surface), **Figure 3A**. Electrodes were lowered into the same coordinates for ACC as the viral infusions and fixed in place. Stainless steel anchor screws were placed across the skull surface and a ground/reference screw was inserted over the cerebellum. Arrays were secured in place with cyanoacrylate, bone cement (C&B Metabond, Parkell Inc., Edgewood, NY), and dental acrylic (Patterson Dental, St. Paul, MN). Animals were provided with post-operative Carprofen (5 mg/kg, *s*.*c*.; Zoetis, Parsippany, NJ), topical anti-biotic ointment (Water-Jel Technologies, Carlstadt, NJ) at the incision site, and oral antibiotics (Sulfamethoxazole and trimethoprim oral suspension USP, Pharmaceutical Associates, Inc., Greenville, SC) daily for five days.

### Drug treatment

Rats were given intraperitoneal (*i*.*p*.) injections of vehicle (VEH: 95% saline + 5% DMSO) or CNO (3 mg/kg CNO in 95% saline + 5% DMSO) 30 min prior to beginning the behavioral task (Stolyarova, Rakhshan et al. 2019, Hart, Blair et al. 2020). As shown in **Figure 1C**, in the discrimination stage upon reaching 80% correct across two consecutive days without drug, rats performed 4 additional sessions of discrimination learning to test for mastery and successful retention. Sessions 1 and 2 after reaching 80% criterion was performed after a single i.p. CNO injection, and Sessions 3 and 4 after a single i.p. injection of vehicle (VEH). If rats did not continue to reach criterion on Session 4 of VEH, subsequent testing sessions were administered with VEH until the criterion was exceeded after which the reversal stage would commence. Injections of VEH or CNO were administered before every reversal session. Once rats reached 50% correct across two consecutive sessions, the other drug was administered for all subsequent reversal sessions until reaching the 80% criterion. Administration of CNO or VEH for the first half (<50%) and VEH or CNO for the second half (>50%) of reversal sessions was counterbalanced across animals.

### Stimulus-based discrimination and reversal learning

After one week of post-surgery recovery, rats began the stimulus-based discrimination learning task (**Figure 1B**). This task was described in detail previously (Aguirre, Stolyarova et al. 2020, Harris, Aguirre et al. 2021), however, in the present design we assigned deterministic stimulus-reward assignments – correct (100%) vs incorrect (0%). Rats initiated each trial by nosepoking a white graphic stimulus in the center screen (displayed for 40 s). The disappearance of the initiation stimulus was immediately followed by two distinct visual stimuli (‘fan’ vs. ‘marbles’) presented on the left and right side of the touch screen (displayed for 60 s) pseudorandomly on each trial. One stimulus was paired with a sucrose pellet reward 100% of the time while the other stimulus was not rewarded. Trials that were not initiated within 40 s were scored as an *initiation omission*, whereas failure to select a choice stimulus was scored as a *choice omission*. Non-rewarded trials were followed by a 5 s time-out, and all trials including omissions were concluded with a 10 s ITI before commencing the next trial. The overall learning criterion constituted meeting the requirement of consuming 60 or more rewards along with selecting the correct option >80% of the trials within a 60 min session across two consecutive days. As above, upon reaching criterion, rats performed an additional four sessions after receiving an injection of CNO or VEH. Rats would then advance to the reversal phase in which the previously correct stimulus was now the incorrect stimulus. Methods and criterion were identical to those described above. Rats then received injections of VEH or CNO on the first day of reversal, and on reaching 50% correct (similarly across 2 days and with 60 or more rewards collected), drug injections were switched until reaching the 80% criterion. Injections of VEH or CNO during the reversal stage was counterbalanced across animals.

### Electrophysiological Recordings

Local field potential (LFP) data in ACC and OFC were acquired at 30 kHz and down sampled to 1,000 Hz. LFPs were bandpass filtered for theta (5-10 Hz) using the *filtfilt()* function in MATLAB. ACC DREADDs placement and electrodes placement are shown in **Figure 3**. Four behavioral task epochs (e.g., trial initiation, correct choice, incorrect choice, and reward port entry) were configured as triggered event outputs from each operant chamber to the data acquisition system that were simultaneously collected with the electrophysiological recordings. Specific events were averaged over trials and over sessions, and subsequently combined across rats.

Data were acquired from a multi-channel data acquisition system (Blackrock Microsystems, LLC., Salt Lake City, UT). The digitizing headstage was connected to the electrode array, and signal was sent to the recording system through a tether and commutator (Dragonfly Inc. Ridgeley, WV). Recording sessions were conducted on all behavioral sessions preceded by an injection. Neural recordings began 15 min after drug injection followed by a 15 min baseline while the animal was inside the operant chamber. The behavioral task commenced at the end of baseline, a total of 30 min post-injection. Recording data for EGFP control animals for all trial epochs are shown in **Figure S3**.

### Histology

At the conclusion of the experiment, rats were anesthetized and administered electrolytic lesions at the recording sites via direct current stimulation (20 μA for 20 s), **Figure 3A**. Rats were euthanized 3 days later by sodium pentobarbital (Euthasol, 0.8 mL, *i*.*p*.; Virbac, Fort Worth, TX) and brains were extracted via transcardial perfusion with phosphate buffered saline (PBS) followed by 10% buffered formalin acetate and post-fixed in this solution for 24 hours followed by 30% sucrose cryoprotection. Tissue was prepared in 40-μM thick coronal sections and either Nissl stained for verification of electrode placement (**Figure 3B**) or cover slipped with DAPI mounting medium (Prolong gold, Invitrogen, Carlsbad, CA) and amplified with NMDAR1 Polyclonal Antibodies (Invitrogen for Thermo Fischer Scientific) for DREADDs verification, visualized using a BZ-X710 microscope (Keyence, Itasca, IL), and analyzed with BZ-X Viewer software. DREADDs expression (visualized by magenta fluorescence) was determined by matching histological sections to a standard rat brain atlas (Paxinos and Watson 2007), **Figure 3C**.

### c-fos immunohistochemistry

Forty μm coronal sections containing ACC were first incubated overnight (16-18 hours) at 4°C in solution containing primary anti-cfos antibody (Anti-cfos (rabbit), 1:500, Abcam, Cambridge, MA, Catalog Number: ab209794), 10% normal goat serum (Abcam, Cambridge, MA, Cat. # ab7481), and 0.5% Triton-X (Sigma, St. Louis, MO, Cat. # T8787) in 1X PBS, followed by three 10-min washes in PBS. The tissue was then incubated for 4 hours in solution containing 1X PBS, Triton-X and a secondary antibody (Goat anti-Rabbit IgG (H+L), Alexa Fluor® 488 conjugate, 1:400, Fisher Scientific, Catalog #A-11034), followed by three 10-min washes in PBS. Slides were subsequently mounted and cover-slipped with DAPI mounting medium (Prolong gold, Invitrogen, Carlsbad, CA), visualized using a BZ-X710 microscope (Keyence, Itasca, IL), and analyzed with BZ-X Viewer software. To verify DREADD-mediated inhibition of neurons in ACC we compared the number of c-fos positive cells in the hM4Di-expressing regions to the number of c-fos positive cells in neighboring (non-hM4Di-expressing, DAPI) areas following vehicle and CNO administration (**Figure 3DE**). Four coronal sections per condition were obtained: hM4Di+VEH, hM4Di+CNO, DAPI+VEH, and DAPI+CNO. There were fewer c-fos positive cells in hM4Di-expressing cells following CNO compared to VEH (*F*(1,5)=8.18, *p*=0.049), but not in DAPI-expressing cells (*F*(1,5)=0.02, *p*=0.903). The difference in c-fos positive cells was not due to sampling differences: the hM4Di-expressing area was not significantly different from the non-hM4Di-expressing (DAPI) area (*t*(10)=-1.536, *p*=0.156).

### Spectral power and Coherence

Local field potential (LFP) data were acquired at 30 kHz and downsampled to 1,000 Hz. Traces were analyzed for signal artifacts through visual inspection and automatic artifact detection in MATLAB. LFP signals that exceeded absolute value of 1.5 mV or when summed cross-band power (2 – 120 Hz) exceeded the 99.98^th^ percentile, were identified as artifact and excluded from all analyses.

Spectral power across frequency bands were determined through a fast Fourier transform via the *spectrogram()* function in MATLAB [frequency bin = 0.5 Hz, 10 s Hanning window]. LFP Coherence (magnitude-squared coherence) was calculated using Welch’s averaged modified periodogram method with a 1-s window and a frequency resolution of 1 Hz via MATLAB function *mscohere()*. LFPs were band-pass filtered for theta (5-10 Hz) using the *filtfilt()* function in MATLAB and designed using the *designfilt()* function (stop bands: 2 Hz, 15Hz; pass bands: 5Hz, 10Hz). Band-passed theta traces were synchronized from −2 to +2 seconds to each of the four behavioral task triggered events (see Data Analysis) in preparation for analyses. Once aligned, the baseline mean (−200 to 0 ms period preceding the triggered event) was subtracted from time=0 to +2 seconds post-event for normalization. Normalized theta for accuracy trail epochs is shown in **Figure S2** and **Figure S3**. ACC-OFC coherence for discrimination and reversal learning is shown in **Figure S5** and **Figure S6**.

### Data analysis

All behavioral and neurophysiological analyses were performed via custom-written code in MATLAB (MathWorks, Inc., Natick, MA). We first demarcated 4 unique task triggered events for our analyses: 1) trial initiation (center stimulus) nosepoke, 2) nosepoke to S+ (correct stimulus or action), 3) nosepoke to S-(incorrect stimulus or action), and 4) reward collection (food magazine head entry).

Learning and performance (speed/latency) data were analyzed with a series of mixed-effects General Linear Models (GLMs) (*fitglme* function; Statistics and Machine Learning Toolbox) first in omnibus analyses that included all factors and both learning phases (discrimination and reversal). For sessions to reach criterion in initial visual discrimination (when animals were not administered drug or recorded), we conducted an independent samples t-test to assess viral group differences. For theta power analyses, 2 × 2 ANOVAs were conducted on baseline-subtracted ACC and OFC theta power with drug (CNO, VEH) x trial type (correct, incorrect) as factors on data averaged from each session. Analyses using trial phase in this case was not treated as a within-subject variable as drug experience differed in each phase: animals received the same order of drug in discrimination (CNO first, then VEH), but drug order was counterbalanced in the reversal phase. Then, each learning phase was analyzed separately for measures where significant interactions of phase were observed. All post-hoc tests were corrected for the number of comparisons (Bonferroni). Statistical significance was noted when p-values were less than 0.05. Major dependent variables included: percent correct (and correct and incorrect choices, probed separately) and median latencies (to initiate a trial, to make the correct choice, to make the incorrect choice, and to collect reward). Pearson correlations for latency measures and theta power were generated using the *corrcoef* function in MATLAB. Latency datapoints exceeding 2 SD were excluded. For latency values, time at trial event (t=0) was subtracted from the previous trial event (t-1). For example, correct or incorrect choice speed was calculated by subtracting time of choice from time of trial initiation, and reward collection speed was calculated by subtracting time of reward collection from time of correct or incorrect choice, in seconds (sec).

To confirm DREADDs efficacy, the number of c-fos positive cells by different drug conditions (CNO, VEH) in hM4Di-expressing cells and in non-hM4Di expressing cells were analyzed with one-way ANOVAs. To ensure that c-fos positive cells differences were not due to differences in sampling of tissue during microscopy, the area of hM4Di spread of virus was compared to that of non-hM4Di (DAPI) and analyzed using an independent samples t-test.

## Acknowledgements

This work was supported by UCLA’s Division of Life Sciences Retention fund (Izquierdo), R01 DA047870 (Izquierdo and Soltani), R21 MH122800 (Izquierdo and Blair), the Training program in Neurotechnology Translation T32 NS115753 (Ye). We appreciate early and helpful comments from members of the Izquierdo lab on these data. We acknowledge the Staglin Center for Brain and Behavioral Health for additional support related to fluorescence microscopy. We thank Alexandra Stolyarova for help with immunohistochemistry. We also thank the NIDA Drug Supply program for the supply of clozapine-N-oxide.

## Conflict of Interest

The authors declare no competing financial interests

## Notes

### Competing Interest Statement

The authors have declared no competing interest.

### Summary of Updates

Manuscript substantially revised in scope and content.

