## Supplemental Figures for "Theta oscillations in anterior cingulate cortex and orbitofrontal cortex differentially modulate accuracy and speed in flexible reward learning"

### Supplementary Figures

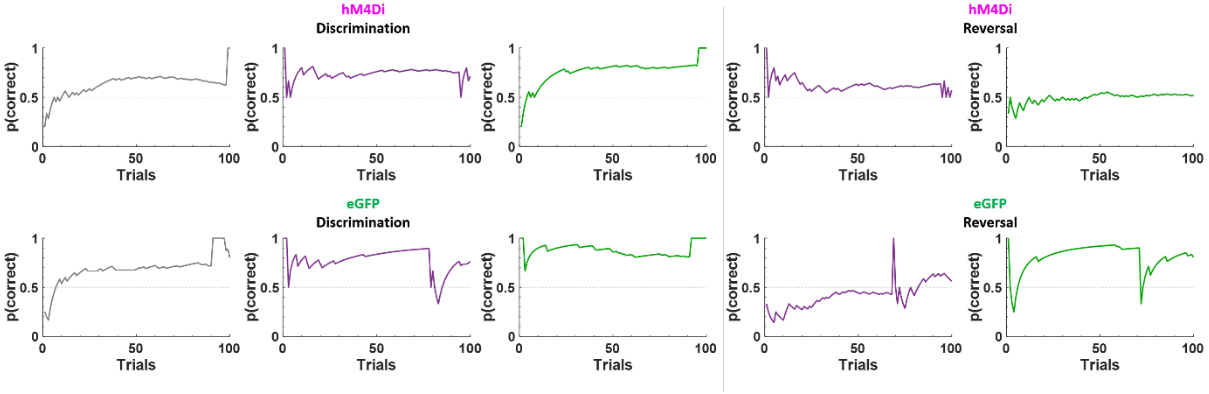


***Figure S1. Trial-by-trial performance within a single session during different learning stages.*** Shown is representative trial-by-trial performance of the first 100 trials within different learning stages for hM4Di (*top*) and eGFP rats (*bottom*). Learning during initial discrimination and reversal for no recording days (grey), and sessions preceded by clozapine-N-oxide (CNO, purple) or vehicle (VEH, green) injections.

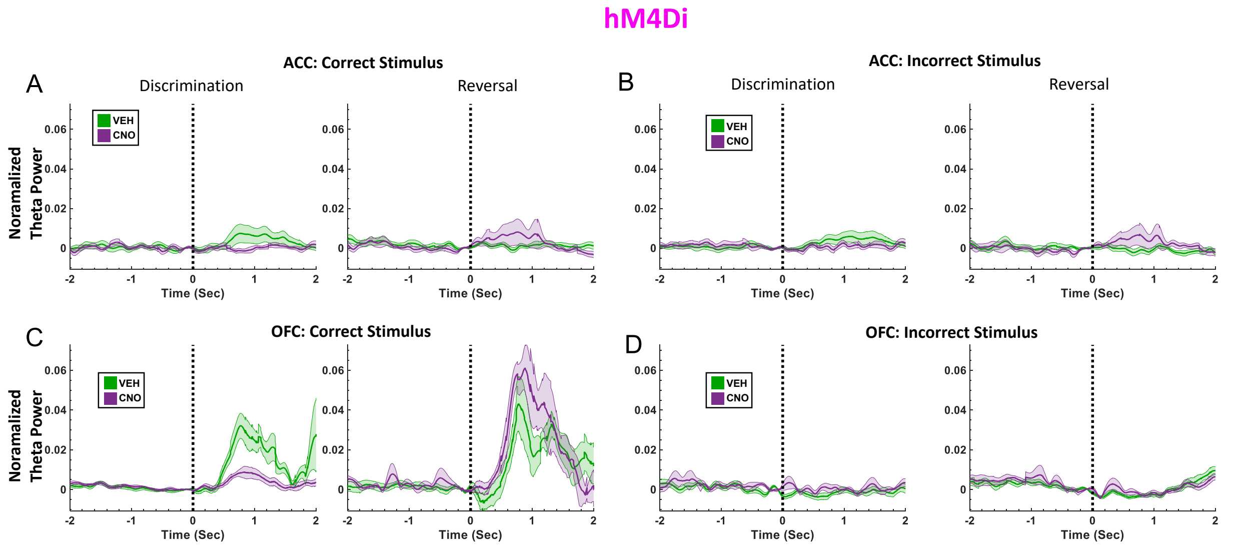


***Figure S2. Normalized theta power during significant trial epochs in learning in hM4Di rats.*** Animals that were prepared with inhibitory Gi DREADDs in ACC were assessed for accuracy in learning that included choice of correct stimulus and choice of incorrect stimulus. (**A**-**B**) Normalized ACC theta power during choice of the correct stimulus and choice of incorrect stimulus during discrimination (*left*) and reversal learning (*right*) phases. (**C**-**D**) OFC theta power during choice of the correct stimulus and choice of incorrect stimulus during discrimination (*left*) and reversal learning (*right*) phases. Animals were injected with clozapine-N-oxide (CNO, purple) or vehicle (VEH, green) 30 min before testing. Theta power mean ± standard error of the mean (S.E.M.) shading.

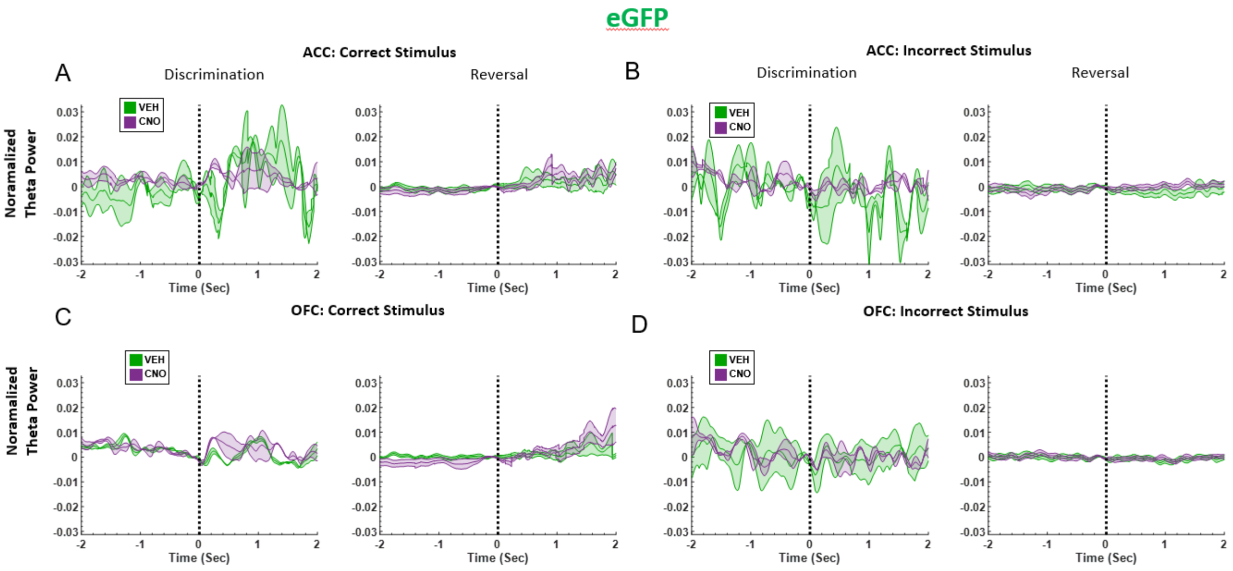


***Figure S3. Normalized theta power during significant trial epochs in learning in eGFP rats.*** Animals that were prepared with eGFP in ACC were assessed for accuracy in learning that included choice of correct stimulus and choice of incorrect stimulus. (**A-B**) Normalized ACC theta power during choice of the correct stimulus and choice of incorrect stimulus during discrimination (*left*) and reversal learning (*right*) phases. (**C-D**) OFC theta power during choice of the correct stimulus and choice of incorrect stimulus during discrimination (*left*) and reversal learning (*right*) phases. Animals were injected with clozapine-N-oxide (CNO, purple) or vehicle (VEH, green) 30 min before testing. Theta power mean ± standard error of the mean (S.E.M.) shading.

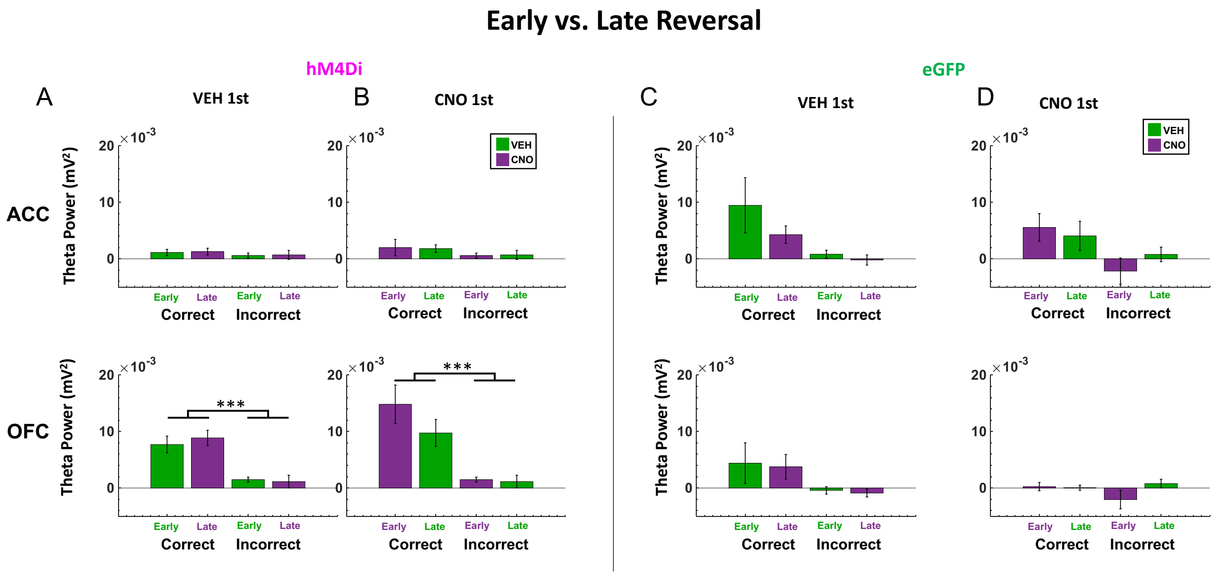


***Figure S4. Average normalized theta power during early and late reversal.*** Baseline-subtracted (normalized) theta power for correct and incorrect choices in the reversal learning phase (data from Figure 4, further separated by early vs. late reversal). (**A**) hM4Di rats were divided into two groups (n=2 each), one beginning the reversal learning phase with administration of VEH (early reversal; green bar). Upon reaching 50% correct criterion, these animals continued each session with administration of CNO instead (late reversal; purple bar). (**B**) The other subset of animals began the reversal phase with administration of CNO and switched to VEH at 50% correct criterion. (**C-D**) As in A and B but for eGFP animals. Patterns were similar to Figure 4, with attenuated ACC theta signal of accuracy, and intact OFC theta signal of accuracy in hM4Di group. Means ± standard error of the mean (S.E.M.) bars. ***p<0.01 following Bonferroni-corrected post-hoc comparisons.

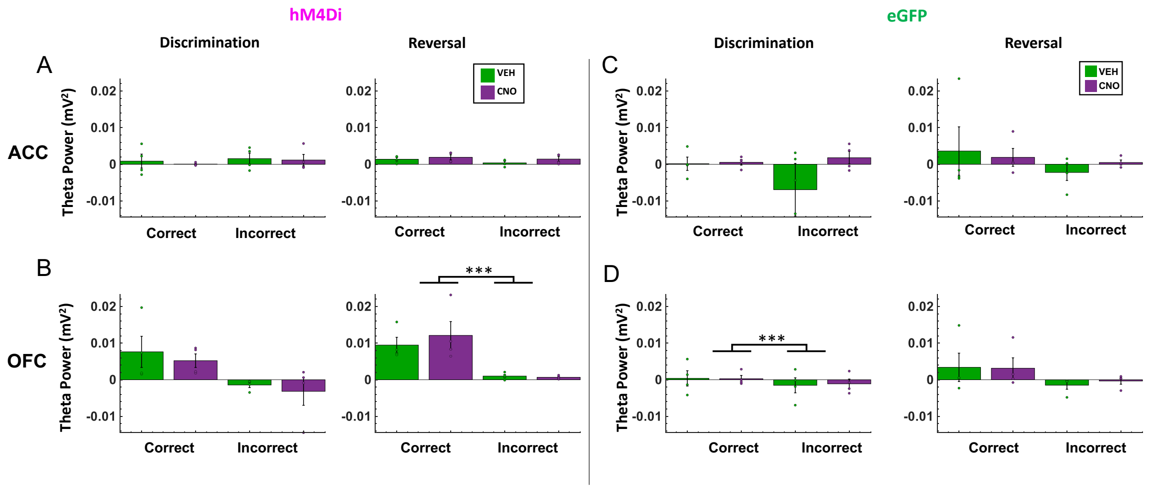


***Figure S5. Individual subject changes in theta power in reversal phase trial accuracy following ACC inhibition.*** Baseline-subtracted (normalized) theta power was calculated for correct and incorrect choices in the discrimination and reversal learning phases (one value per animal, per condition). (**A**) In hM4Di animals, there was no significant difference in normalized ACC theta power between correct vs. incorrect trials in the reversal phase. (**B**) In hM4Di animals, there was no significant difference in normalized theta power in OFC for correct vs. incorrect trials in the discrimination phase, but there was significant OFC theta signaling trial accuracy in reversal learning. (**C)** In eGFP animals, we observed no significant difference in normalized theta power in ACC for correct vs. incorrect trials in discrimination or reversal. (**D**) In eGFP animals, normalized theta power in OFC signaled trial accuracy in the discrimination, but not in the reversal, phase. Patterns were similar to Figure 4, with attenuated ACC theta signal of accuracy, and intact OFC theta signal of accuracy in the hM4Di group. Means as individual scatter plots ± standard error of the mean (S.E.M.) bars. ***p<0.01 following Bonferroni-corrected post-hoc comparisons.

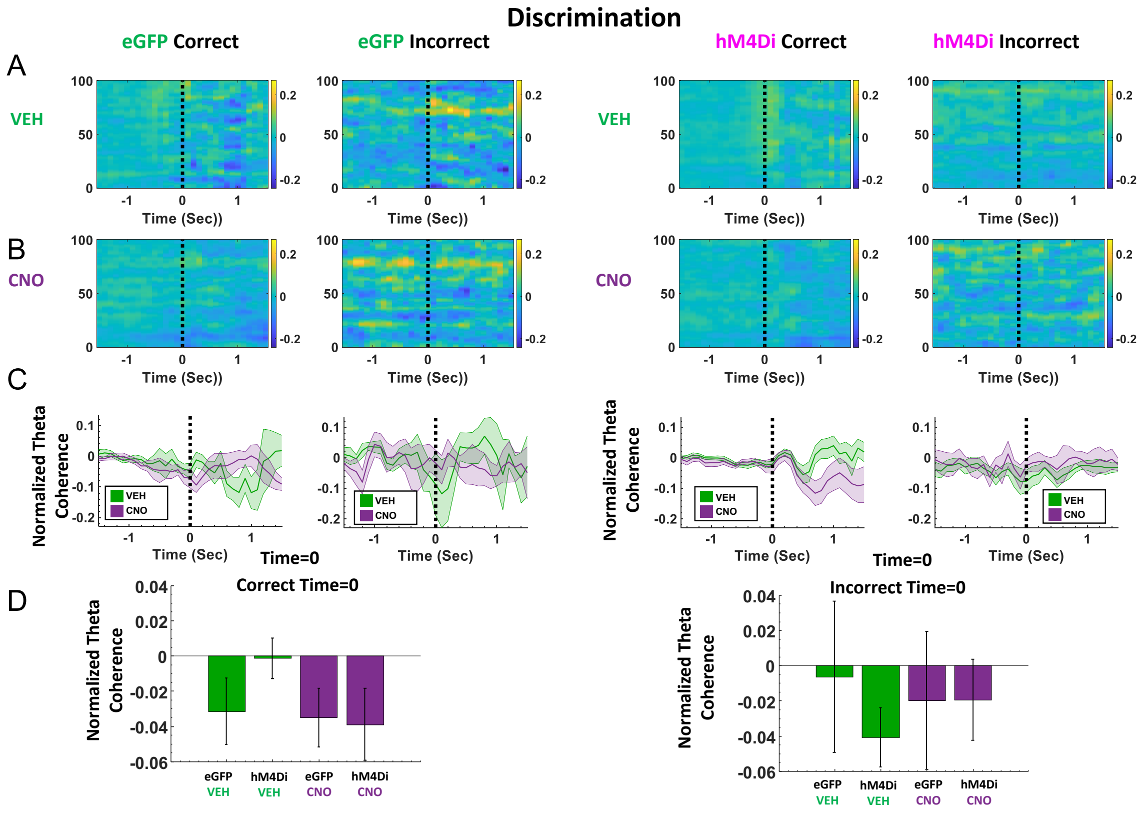


***Figure S6. ACC-OFC theta coherence during discrimination trial epochs.*** Local field potentials were aligned to the choice of correct and incorrect stimulus (time=0) of discrimination trials, the average baseline power (-200 to 0 ms period preceding the choice) was subtracted from the entire -2 to +2 period. Average normalized spectral coherence for eGFP and hM4Di animals in the VEH (**A**) and CNO (**B**) conditions. (**C**) Isolated theta coherence from A and B comparing VEH and CNO conditions. (**D**) Quantified normalized theta power at time=0 of the choice epoch, A 2x2 ANOVA (drug x trial type) did not reveal any statistically significant differences in theta coherence. Mean coherence ± standard error of the mean (S.E.M.) shading and bars.

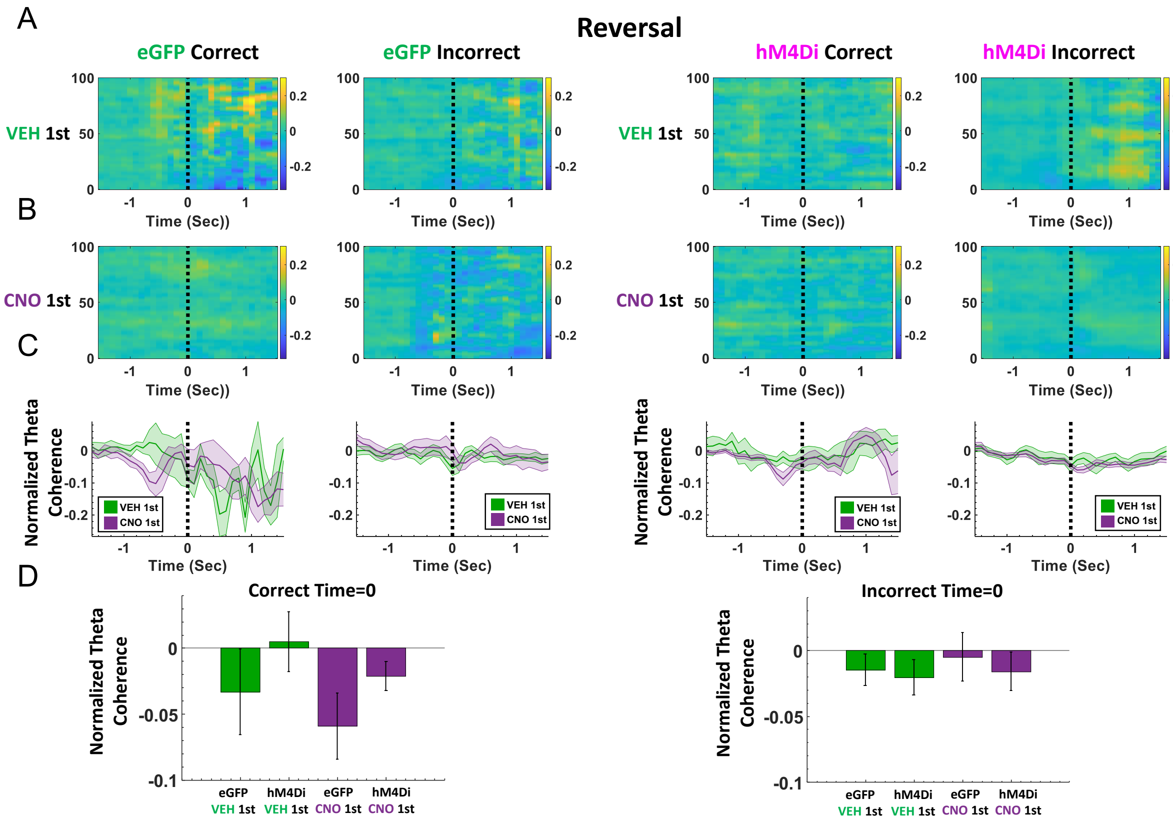


***Figure S7. ACC-OFC theta coherence during reversal trial epochs.*** Local field potentials were aligned to the choice of correct and incorrect stimulus (time=0) of reversal trials, the average baseline power (-200 to 0 ms period preceding the choice) was subtracted from the entire -2 to +2 period. Average normalized spectral coherence for eGFP and hM4Di animals in the VEH **(A)** and CNO **(B)** conditions. **(C)** Isolated theta coherence from A and B comparing VEH and CNO conditions. **(D)** Quantified normalized theta power at time=0 of choice epoch, a 2x2 (drug x trial type) ANOVA did not reveal any statistically significant differences in theta coherence. Mean coherence ± standard error of the mean (S.E.M.) shading and bars.
